# Genotyping and functional regression trees reveals environmental preferences of toxic cyanobacteria (*Microcystis aeruginosa* complex) along a wide spatial gradient

**DOI:** 10.1101/2019.12.20.885111

**Authors:** Gabriela Martínez de la Escalera, Angel M. Segura, Carla Kruk, Badih Ghattas, Claudia Piccini

## Abstract

Addressing ecological and evolutionary processes explaining biodiversity patterns is essential to identify the mechanisms driving community assembly. In the case of bacteria, the formation of new ecologically distinct populations or ecotypes is proposed as one of the main drivers of diversification. New ecotypes arise when mutation in key functional genes or acquisition of new metabolic pathways by horizontal gene transfer allow the population to exploit new resources, making possible their coexistence with parental population. Recently, we have reported the presence of toxic, microcystin-producing organisms from the *Microcystis aeruginosa* complex (MAC) through a wide environmental gradient (800 km) in South America, ranging from freshwater to estuarine-marine waters. In order to explain this finding, we hypothesize that the success of toxic organisms of MAC in such array of environmental conditions is due to the existence of ecotypes having different environmental preferences. So, we analysed the genetic diversity of microcystin-producing populations of *Microcystis aeruginosa* complex (MAC) by qPCR and high resolution melting analysis (HRMA) of a functional gene (*mcyJ*, involved in microcystin synthesis) and explored its relationship with the environmental conditions through the gradient by functional classification and regression trees (*f*CART). Six groups of *mcyJ* genotypes were distinguished and selected by different combinations of water temperature, conductivity and turbidity, determining the environmental preferences of each group. Since these groups were based on the basis of similar sequence and ecological characteristics they were defined as ecotypes of toxic MAC. Taking into account that the role of microcystins in MAC biology and ecology has not yet been elucidated, we propose that the toxin might have a role in MAC fitness that would be mainly controlled by the physical environment in a way such that the ecotypes that thrive in the riverine zone of the gradient would be more stable and less influenced by salinity fluctuations than those living at the marine limit of the estuary. These would periodically disappear or being eliminated by salinity increases, depending on the estuary dynamics. Thus, ecotypes generation would be an important mechanism allowing toxic MAC adapting to and succeed in a wide array of environmental conditions.

## 1. Introduction

Microbial communities are composed by a myriad of genetic variants that interact and respond rapidly to changing environments (Koeppel et al., 2008). Deep ecological and physiological differences among microorganisms’ populations mirror differences at the genomic level (Welch et al., 2002). These differences are usually difficult to detect when using slowly evolving phylogenetic markers such as ribosomal genes, which circumvents the elucidation of the mechanisms accounting for the observed community structure. Therefore, developing a theoretical framework and appropriate tools to quantify and understand the environmental forces shaping community structure is timely relevant. One of these frameworks is the ecotype theory of bacterial species (Cohan, 2002, 1994; Cohan and Perry, 2007; Ward et al., 1994), which defines the ecotype as a clade of phylogenetically-related microorganisms (meaning that they belong to the same bacterial species) sharing ecological characteristics. An ecotype generates after a single individual experiences mutation or recombination that changes its ecology, allowing the utilization of a new set of resources or to thrive under a particular environmental condition (Cohan, 2002). Under this framework, it has been defined that two individuals belong to the same ecotype when their 16S rRNA genes share a high degree of identity (97 - 99%) (Stackebrandt, 2006) and ecological preferences (Kopac and Cohan, 2011). Depending on the mechanism that induces the variation, different models of ecotype generation have been proposed (Cohan, 2006; Cohan and Ward, 2005; Gevers et al., 2005).

In particular, ecotypes have been defined for several cyanobacteria species, such as *Prochlorococcus* (Moore, 1998), *Synechococcus* (Sohm et al., 2016) and *Cylindrospermopsis* (Piccini et al., 2011). Different ecotypes have been described for these organisms based on their environmental preferences (e.g. temperature, light intensity, nutrients concentrations and iron availability) (e.g. size, shape) and physiology (e.g. nitrogen fixation). As an ecological strategy, the existence of ecotypes adapted to different temperatures (Chandler et al., 2016) and light intensities (Moore et al., 1995) allowed *Prochlorococcus* to account for the 50%of the total chlorophyll and an estimated production of 4 gigatons carbon fixed per year in vast areas of the surface ocean (Biller et al., 2015), allowing this microorganisms to occupy the entire euphotic zone. There are several examples in the literature identifying ecotype diversification as one of the most relevant mechanisms explaining the colonization of different potential niches in many environments, and is proposed as a main driver of bacterial speciation (Cohan, 2007).

Several methods and algorithms have been applied to identify bacterial ecotypes and to address their presence and abundance, which are usually related to PCR amplification of phylogenetic marker genes and posterior analysis of the amplicons, either by fingerprinting methods (e.g. DGGE, TGGE) (Ferris et al., 2003) or by sequencing (Chandler et al., 2016; Martiny et al., 2009). Fingerprinting-based methods are described to discern between sequences differing in a single nucleotide, while in the case of sequencing (usually 16S genes or the internal transcribed spacer of the bacterial ribosomal operon) a similarity cut-off is needed. This cut-off can produce different results, depending on how broadly the taxa are defined. Thus, identifying ecotypes by this approach would depend on the stringency of taxa assignment (Martiny et al., 2009). In the case of High Resolution Melting Analysis (HRMA), which is a fingerprinting-based method, it has been used to study diversity at the genotype level (Hjelmsø et al., 2014; Kim and Lee, 2014). The HRMA denaturation curves give information about the whole amplicon sequence, detecting single nucleotide polymorphisms (SNPs) with great precision at high throughput and relatively low cost. Therefore, it provides melting profiles which curves are used to discriminate among different genetic populations. HRMA has been applied to genotyping, mutation scanning and SNPs detection in bacterial populations (Hofinger et al., 2009; Thomsen et al., 2012; Tong and Giffard, 2012; Wittwer et al., 2003) and human disease (Li et al., 2011) as well as to address diversity of microbial communities (Hjelmsø et al., 2014; Kim and Lee, 2014; Zeyoudi et al., 2015). The output of HRMA denaturation curves present high dimensionality and correlation that can be managed by machine learning techniques, providing an efficient way to interpret the information (Libbrecht and Noble, 2015). In particular, functional analysis allows an objective analysis of correlated results, thus, the combination of molecular methods with machine learning techniques appears as a promising way to explore microbial community assembly patterns under the ecotype theoretical framework.

In the case or *Microcystis* genus of cyanobacteria, several morpho-species sharing nearly identical 16S rDNA gene sequence (more than 97% identity) but exhibiting ecological distinctness have been described (Otsuka et al., 1998). *Microcystis* is one of the most common bloom-forming cyanobacterial genus worldwide, capable of successful growth in a variety of freshwater and brackish ecosystems (De Leon and Yunes, 2001; González-Piana et al., 2017, 2011; O’Neil et al., 2012; Srivastava et al., 2013). It harbours a number of species defined according to their morphology and ecology that are aggregated in the *Microcystis aeruginosa* Complex (MAC), whose diversity and bloom formation capacity varied along environmental gradients from headwaters to marine waters (Kruk et al., 2017; Martínez de la Escalera et al., 2017; Sabart et al., 2009; Segura et al., 2017; Tanabe et al., 2018). Most of MAC organisms are able to produce microcystin, a toxin known to cause of serious liver diseases in humans (Azevedo et al., 2002; Dittmann and Wiegand, 2006; Milutinović et al., 2003; Vidal et al., 2017) with more than 250 variants (Meriluoto et al., 2017; Puddick et al., 2014), which are produced via non-ribosomal peptide synthesis (NRPS) and polyketide synthase (PKS) (Dittmann et al., 1997; Tillett et al., 2000).

MAC blooms can consist of mixtures of microcystin-producing (toxic genotypes) and non-producing (non-toxic genotypes) populations (Kaebernick and Neilan, 2001; Kurmayer and Kutzenberger, 2003; Vezie et al., 1998). There is no consensus about the role of microcystins in the biology and ecology of MAC organisms. Environmental factors, such as nutrients, salinity, light intensity and temperature have been described as influencing the toxin concentration (Dittmann et al., 1997; Kameyama et al., 2004; Lee et al., 2000). Recently, Qu et al. (2018) reported for *M. aeruginosa* isolates that metabolic functions essential for cell growth (photosynthesis efficiency, carbohydrate metabolism and redox homeostasis) are important in microcystin production. Furthermore, it has been proposed that, under oxidative stress conditions, microcystin increases the fitness of *Microcystis* by modulating a number of proteins (Zilliges et al., 2011). Owing to their lifestyle, cyanobacteria of MAC are exposed to very high irradiances and oxygen over-saturation, especially during blooms (Zilliges et al., 2011). So, the presence of microcystin would help organisms survival. If microcystin presence is important for the fitness of the organisms and its production is regulated by specific environmental conditions, we hypothesized that the acquisition and posterior modification of the *mcy* genetic cluster could drive the ecological diversification of MAC.

The morphological and ecological differences used to define *Microcystis* species are not reflected in the 16S rDNA-based phylogeny, so different theoretical and practical approaches are needed to define ecologically relevant species. For example, application of multilocus sequence typing using seven housekeeping loci has shown that *M. aeruginosa* is divided into at least seven distinct phylogenetic clusters matching partially the colony morphology and microcystin production (Tanabe et al., 2009). Nonetheless, Pérez-Carrascal et al. (2019) have recently reported a study on *Microcystis* diversity based on the genomic sequencing of several isolates belonging to different species and found that the morpho-species *M. aeruginosa* was in fact distributed across 12 genomic cluster, which indicate that is not a coherent species and that most of the ecological information gathered for this organism should be considered as belonging to *Microcystis* genus. Thus, MAC offers an excellent model group to explore microbial community assembly based on genetic variability using the ecotype framework.

Here, we used MAC communities from a large environmental gradient (ca. 800 km) as a model to evaluate if the ability of these organisms to thrive in a wide range of environments is based on the existence of multiple genotypes adapted to different sets of environmental conditions (ecotypes). To identify ecotypes, MAC genotyping was based on *mcyJ* melting curves obtained from HRMA, which were later classified based on environmental variables by a functional classification and regression trees approach (*f*CART).

## 2. Materials and methods

### 2.1. Strategy

High Resolution Melting is post real time-PCR method used to perform genotyping based on the detection of single nucleotide polymorphisms (SNP). After PCR, a melting analysis is performed by gradually heating the amplicons at 0.1 °C steps. During this process, as the temperature increases the melting point of the amplicon is reached and amplicon DNA denatures, melting apart the double strand and making the fluorescence of the dye to fade away. This melting behaviour and concomitant fluorescence decay is represented as melting curves (fluorescence decay during melting temperature increase) and is characteristics for each amplicon sequence. Thus, it can discriminate samples according to their sequence length, GC content and strand complementarity. This is the case when analysing sequences from single isolates, however, our goal was to detect variations between melting curves of *mcyJ* gene from different communities, as a single environmental sample included the DNA of a whole MAC community. Therefore, the resulting melting curve correspond to the abundance-weighted average melting profile of all the *mcyJ* gene sequences (see section 2.7 below for further information). The relationships between the structure of the toxic MAC communities and the environmental variables were evaluated with a functional classification and regression trees (*f*CART) (Breiman et al., 1984; Nerini and Ghattas, 2007). *f*CART is an extension of classical Classification and Regression Tress used in ecology (Breiman et al., 1984; De’ath, 2002; Nerini and Ghattas, 2007) in which the response variable is no longer a given value, but a function (Nerini and Ghattas, 2007). This methodology was used to evaluate the relation of environmental variables and toxic genotypes and identify group of closely-related toxic genotypes of MAC (ecotypes) exhibiting the same environmental preferences.

### 2.2. Study site

The study area is located in the subtropical region of South America and covered an extension of ca. 800 km, from Salto Grande reservoir in the Uruguay river (31° 11’ latitude, 57° 52’ longitude) to Punta del Este (34° 57’latitude, 55° 02’ longitude), at the marine end of Río de la Plata estuary. Six sites were sampled every two months during one-year (from January 2013 to March 2014) and subsurface (~ 0.5 m) samples were taken in coastal stations (0.01-0.5 km) (for more details see (Martínez de la Escalera et al., 2017). In total, 36 water samples were analysed. The system presents strong temporal and spatial gradients in terms of temperature, conductivity (which was used as a proxy of salinity) and turbidity (Acha et al., 2008; Ferrari et al., 2011; García-Rodríguez et al., 2013; Kruk et al., 2015; Martínez de la Escalera et al., 2017). The highest surface water temperatures in Salto Grande reservoir are usually recorded during summer (January to March, 19-33 °C) while the lowest temperatures belong to the outer marine zone of the Río de la Plata during winter-early spring (June to October, 11-18 °C) (Kruk et al., 2017; Martínez de la Escalera et al., 2017). Conductivity was minimum in freshwater systems (0.023 mS cm^−1^, Salto, Fray Bentos, Carmelo and Colonia) and maximum at the estuary (55 mS cm^−1^, Montevideo and Punta del Este) (Kruk et al., 2017; Martínez de la Escalera et al., 2017). Turbidity ranged from 0 to 187 NTU, with higher values at the middle of the gradient (Stns. Carmelo and Colonia) (Kruk et al., 2017; Martínez de la Escalera et al., 2017). MAC biovolume was dominant in Salto Grande reservoir during summer and decreased toward the marine end (Punta del Este) (Kruk et al., 2017; Martínez de la Escalera et al., 2017). The presence of *mcy* genes (*mcyB*, *mcyD*, *mcyE* and *mcyJ*) was detected by qPCR in the whole ecosystem while maximum abundances were detected in summer and spatially decreasing from the reservoir to the marine sites (Martínez de la Escalera et al., 2017).

### 2.3. DNA extraction

For the DNA extraction, 250-300 ml of the subsurface water samples collected with clean plastic 20 L carboys were filtered through 0.22 μm sterile polycarbonate membrane (Millipore, Darmstadt, Germany), which were immediately frozen at −20 ºC until processing. Procedures for nucleic acid extraction were performed as described in (Martínez de la Escalera et al., 2014).

### 2.4. Quantification of mcyJ gene in the environmental samples

Two microliters of DNA extracts from each sample (ca. 50 ng DNA) were applied to the Power SYBR Green PCR (Invitrogen) with a final reaction volume of 20 ml. Primers for mcyJ gene were those from Kim et al. (2010). Cycling conditions were 2 min at 50 °C, 15 min at 95 °C and 40 cycles of 15s at 94 °C, 30s at 60 °C and 30s at 72 °C, including a last melting step from 65 to 95 °C at increases of 1 °C each 4 s. A 96 FLX Touch TM thermal cycler (Bio-Rad) was used. To quantify the abundance of *mcyJ* gene, cloned amplicons (Martínez de la Escalera et al., 2017) were used to perform the calibration curves. Curves were achieved using five serial dilutions from 1/10 to 1/100,000 of the cloned genes (in quintuplicates) and applied to qPCR in the same PCR plate where the samples were assayed. Samples were run in triplicate.

### 2.5. High Resolution Melting Analysis (HRMA) of mcyJ amplicons

Amplification of the *mcyJ* gene was performed using the HRM primers described in the literature (Kim et al., 2010). PCR amplification was conducted using a 96 FLX Touch TM thermal cycler (Bio-Rad, California, USA). Two microliters of DNA extracts from each sample (ca. 50 ng DNA) were applied to the MeltDoctor HRM Master Mix (Applied Biosystems, California, USA) with a final reaction volume of 20 μL. Cycling conditions were 2 min at 50 °C, 15 min at 95 °C and 40 cycles of 15 s at 94 °C, 30 s at 60 °C and 30 s at 72 °C. To obtain the HRM melting profiles, RFU (relative fluorescence units) obtained from each sample within a melting region from 65 °C to 95 °C at 0.02 °C/s increases, were recorded. HRM data were acquired using Bio-Rad Precision Melt Analysis (Bio-Rad, California, USA) and each sample was run in duplicate. All the samples were run in the same PCR plate. Melting curves were normalized to the same fluoresces level (RFU) using the pre- and post-melt regions (before and after the melting region, respectively). Pre- and post-melt region were selected based on the specific melt region of the *mcyJ* amplicon (from 75 ºC to 82 ºC, Tm=79.5 ºC). These relative values of RFU were used for the statistical analysis.

### 2.6. Functional CART

The melting curves (RFU) were first represented in a functional basis using a non-periodic β-splin basis of order 4 (this was the optimal choice after several experiments). The multivariate output CART (De’ath, 2002; Nerini and Ghattas, 2007) was used to model these coefficients (output variables) using the following explanatory variables as input: total nutrients (Total Nitrogen; TN and Total Phosphorus; TP), wind intensity (WI), water temperature (T), turbidity (Turb) and conductivity. Cross-validation was used to optimize the size (number or leaves) of the final tree. To assess the performance (prediction error) and the reliability of such an optimal tree, we repeated the following steps 100 times: i) random split of the data in two parts, learning and test sample (in proportion 2/3, 1/3 respectively); ii) building of an optimal tree over the learning sample; iii) random permutation of the observed input variable and building of another optimal tree over the "permuted" data set; iv) computing prediction error of optimal and permuted trees separately. The error distribution between both trees (in the 100 replicates) was compared using a log-likelihood ratio test (LRT). Finally, we evaluated the structure of the tree with respect to MAC community structure by evaluating the distribution of relevant ecological traits for toxic cyanobacteria (volume, surface/volume ratio, MAC biovolume and richness, and *mcyJ* gene abundance). This was done using LRT and Tukey’s pairwise post-hoc comparisons. All statistical analyses were performed with the free software R, version 3.6.1 using {fda}, {rpart}, {nlme} and {PMCMR} packages (Pinheiro et al., 2014; Pohlert, 2014; R Core Team, 2013; Ramsay et al., 2014; Therneau et al., 2015).

### 2.7. Identification of toxic ecotypes

To explain the basic ideas on how the ecotypes of toxic MAC were identified, we give a synthetic example with the rationale and operative steps of the method. In first place, we assumed that *mcyJ* melting profile obtained from a given MAC community (a sample) represents the abundance-weighted average profile of all the genotypes present in that sample. After running simulations and in agreement with our assumption, the combination of the hypothetical environmental drivers A and B (*e.g.* temperature and salinity) deterministically defined the abundance of 3 particular *mcyJ* genotypes (J1, J2 and J3; Fig 1a), which are expressed by their individual melting profile (Fig 1b). So, under a particular environmental arrangement of A and B, we expect to find a given combination of genotype abundances in the community (Fig 1c). The relative abundance of each genotype under each environmental array of conditions determine a distinctive community melting profile, *i.e.* an ecotype (Fig 1c; grey dashed line). For our simulated community, the relationship between abundance and environmental conditions were constructed from a bivariate normal distributions with average µ_*i*_ ={µ_A_,µ_B_} and covariance matrix σ_*i*_ specified for each genotype, where *i* = 1:3. The three *mcyJ* genotypes used here (J1, J2 and J3) belonged to previously obtained *mcyJ* clones showing different melting curves (Martínez de la Escalera et al., 2017). After obtaining the melting curves from each genotype, we randomly sampled environmental conditions 50 times and, for each condition (defined by the pair {A,B}), the relative abundance of each genotype was estimated and a community melting profile was constructed. Under our hypothesis, each region in the environmental space should be characterized by a similar community melting profile corresponding to a given toxic ecotype (Fig 1d). It is important to recall that environmental drivers can be continuous or categoric and explicitly include biotic interactions among the defining variables. In the extreme case where the abundance of the three *mcyJ* amplicons is not determined by the environmental variables selected (µ_*i*_ = µ_*j*_ and σ_*i*_ = σ_*j*_ for all *i, j*), a single ecotype is expected. Under this situation, the functional CART would not partition the data into separated ecotypes, and the tree will remain as a “root tree”.

**Fig 1.**
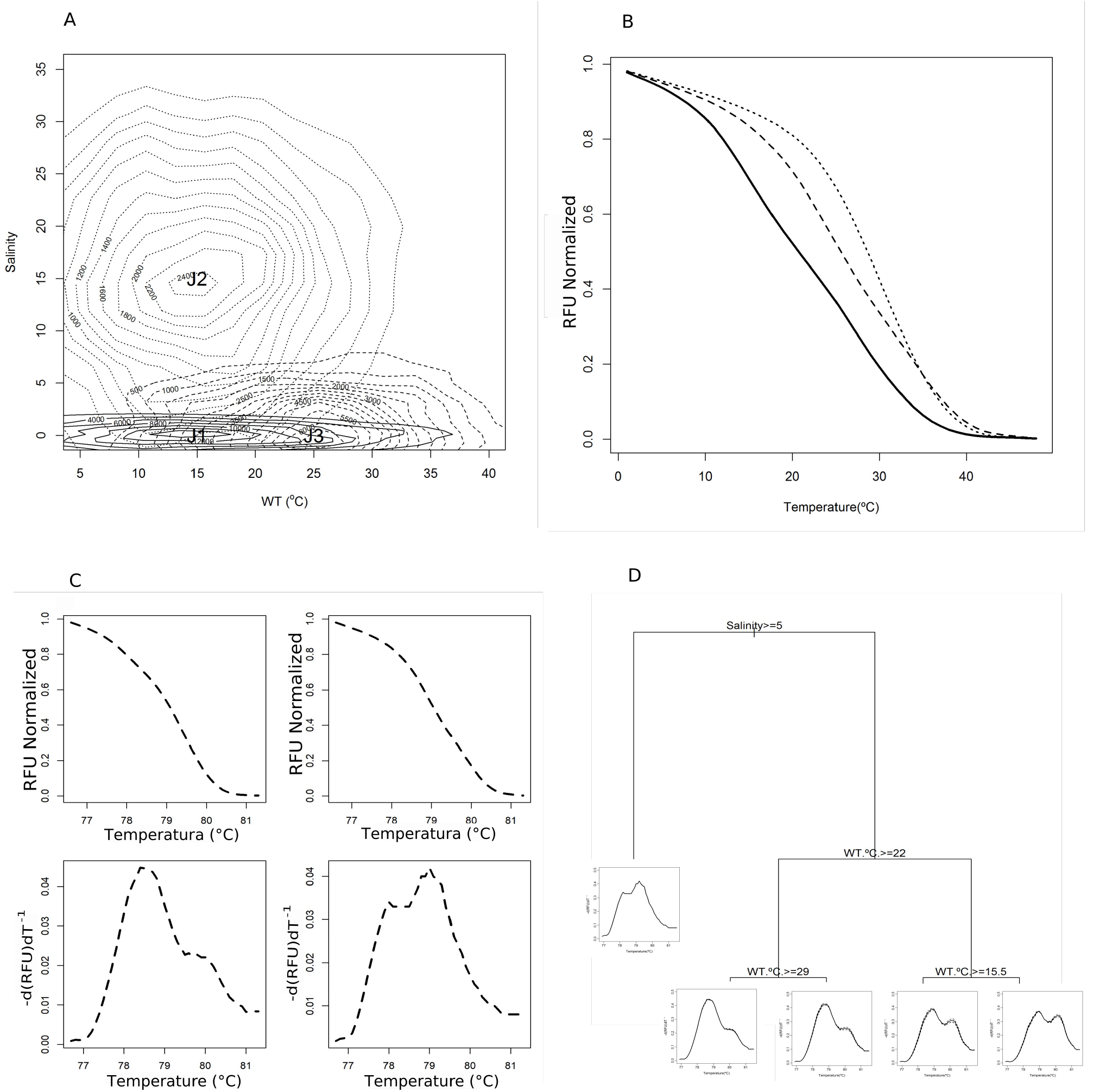
Synthetic example with the rationale and operative steps followed for the HRMA-*f*CART method. A) Abundance and distribution of three *mcyJ* genotypes (J1, J2 and J3) according to environmental characteristics (*e.g.* temperature and salinity). B) Normalised melting curves from the three HRMA curves obtained from three cloned *mcyJ* individually analysed: J1 (solid line), J2 (dashed line) and J3 (dotted line). C) Normalised HRMA curves obtained from a single genotype. Upper left, curve obtained from a genotype thriving in freshwater at 33 °C and its derivative below; upper right, curve obtained from a genotype thriving in marine water (salinity 33) and 13 °C with its derivative below. D) Optimal functional regression tree obtained with randomly sampled environmental conditions. In each node, the environmental variable and its threshold value are shown. Water temperature (WT), and salinity. At the end of each branch the average melting peak (solid line) and its standard deviation (dashed line) representing toxic genotype community are shown.

## 3. Results

The *f*CART analysis of HRMA profiles allowed to summarize MAC community composition and responses along an 800 km environmental gradient into six groups of *mcyJ* genotypes having distinct environmental preferences and that were defined as ecotypes. The ecotypes (named from A to F) were detected using 72 normalized RFU melting profiles and three environmental variables selected as relevant: temperature, salinity and turbidity (Fig 2; Table 1). The optimal tree had an average prediction error significantly lower than the average error obtained from permuted trees (LRT, p<0.05; Fig 3).

**Table 1.** Environmental variables associated to each Ecotype of toxic MAC. Mean values and ranges of environmental variables associated to each ecotype: temperature (Temp, ºC), conductivity (Cond, mScm^−1^), turbidity (Tur, NTU), *mcyJ* gene abundance (copies ml^−1^), and the relative frequency of occurrence of each ecotype. BDL = below detection limit.

| Ecotype | Niche | Temp (°C) | Zone | Cond (mScm<sup>-1</sup>) | Tur (NTU) | *mcyJ* gene (copies ml<sup>-1</sup>) | Ecotype relative frequency of occurrence |
| --- | --- | --- | --- | --- | --- | --- | --- |
| A | Warm brackish water | 19.1 (14.8-22.9) | Middle and outer estuary | 36.4 (18.2-52.0) | 40.9 (15.4-89.6) | 34.9 (BDL-96.0) | 0.14 |
| B | Hot freshwater | 27.2 (24.3-33.6) | Reservoir and riverine freshwater | 0.041 (0.023-0.054) | 28.5 (14.3-47.9) | 2.30E4 (BDL-1.14E5) | 0.14 |
| C | Warm freshwater | 20.5 (15.6-23.4) | Reservoir and riverine freshwater | 0.074 (0.033-0.113) | 56.0 (19.9-127.0) | 1.21E4 (BDL-4.54E4) | 0.31 |
| D | Warm water and low turbidity | 20.9 (16.1-25.9) | Inner and middle estuary | 14.02 (0.048-52.7) | 7.6 (BDL-14.2) | 46.2 (BDL-148.7) | 0.20 |
| E | Cold brackish water | 11.9 (11.2-12.3) | Middle and outer estuary | 29.13 (0.113-55.8) | 20.8 (BDL-49.0) | 11.7 (BDL-44.7) | 0.11 |
| F | Cold freshwater | 12.7 (11.0-14.2) | Reservoir and riverine freshwater | 0.061 (0.036-0.095) | 17.5 (0.9-42.2) | 192.7 (0.3-437.2) | 0.09 |

**Fig 2.**
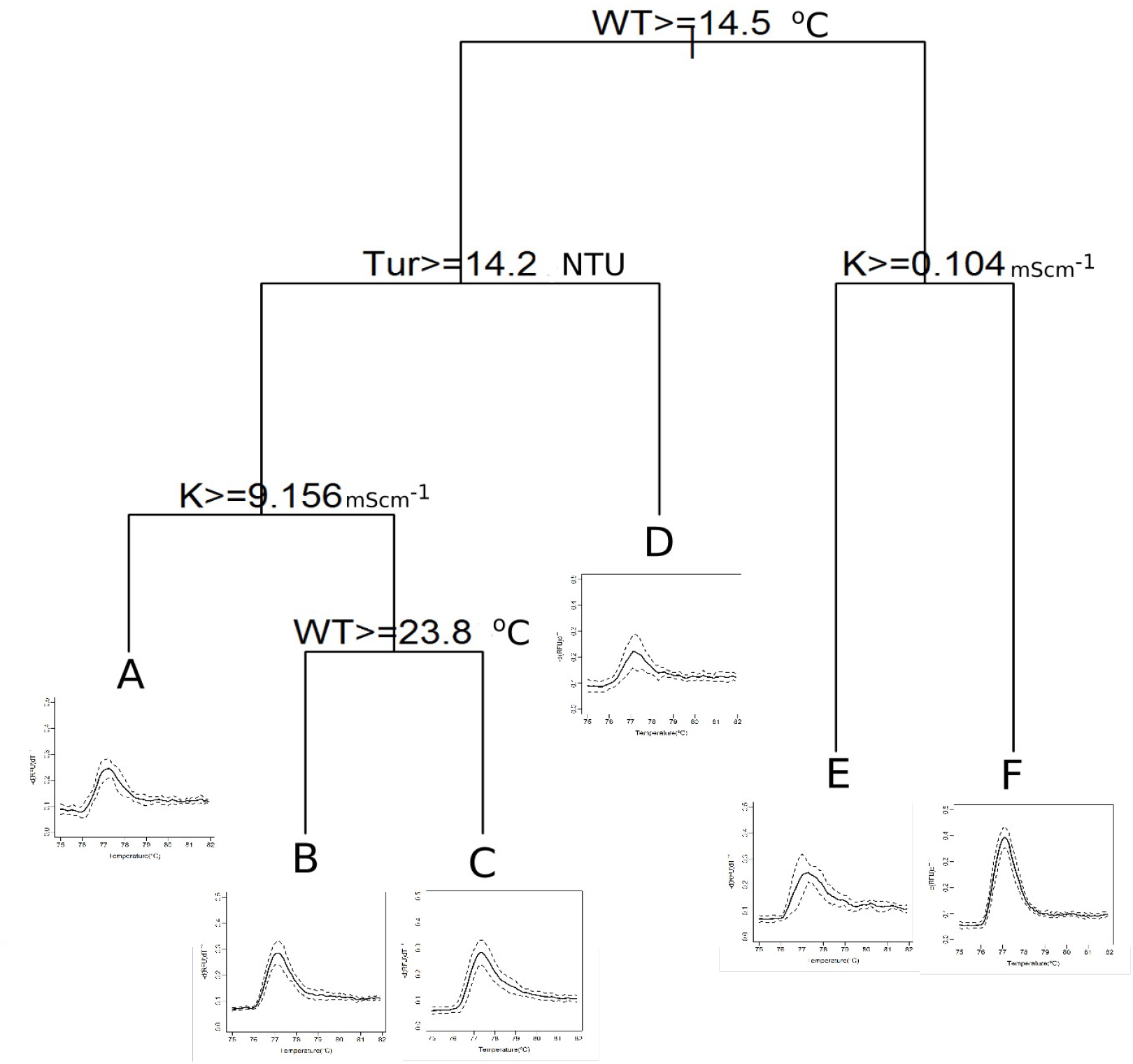
Optimal functional regression tree. Functional regression tree showing the main environmental variables explaining the profile diversity of toxic genotypes. In each node, the environmental variable and its threshold value are shown. Water temperature (WT), turbidity (Tur) and conductivity (K). At the end of each branch the average melting peak (solid line) and its standard deviation (dashed line) representing toxic genotype community are shown.

**Fig 3.**
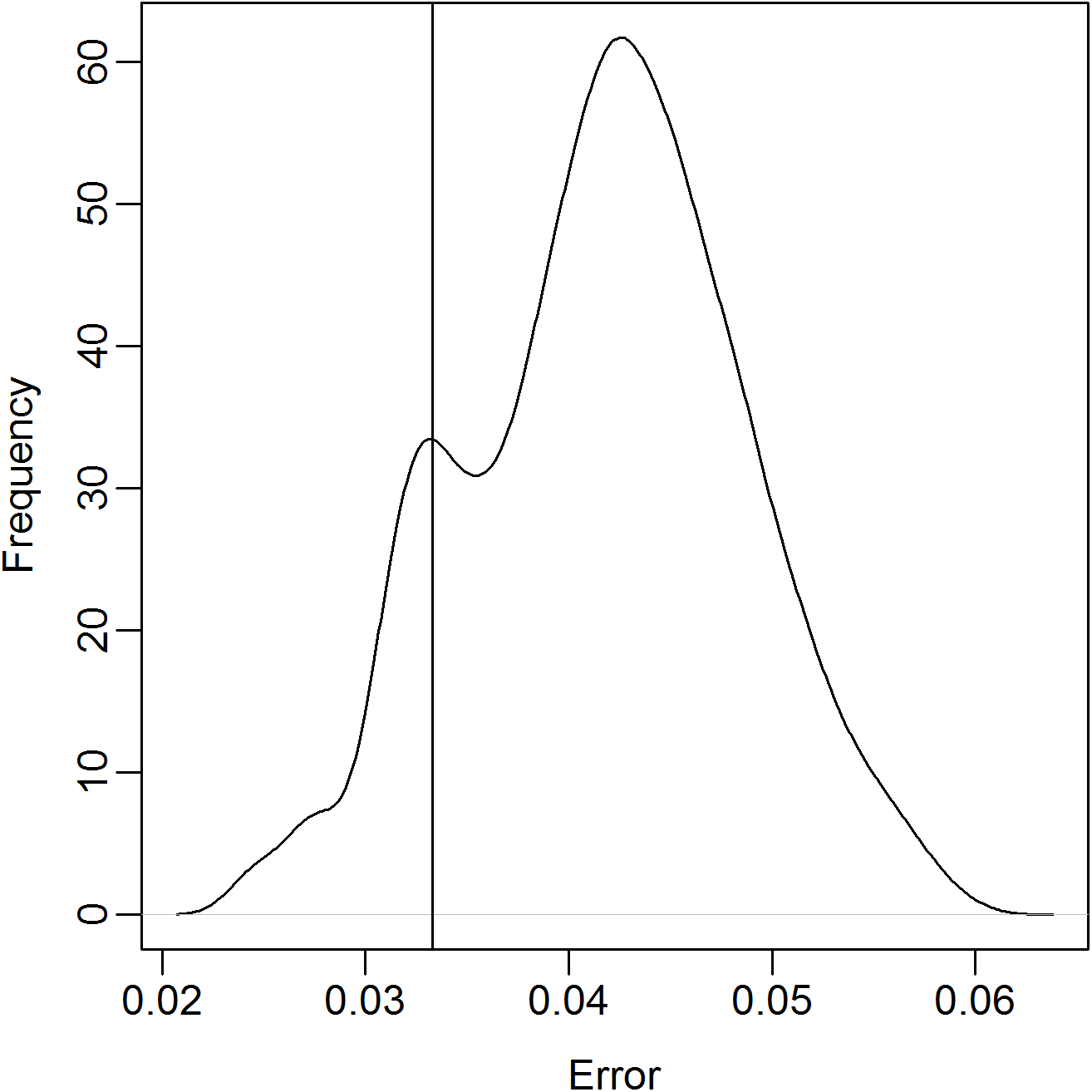
Errors obtained for functional regression trees. Density of the tree-error performed by randomly switching the values of environmental variables and mean value of the error obtained using the original data.

Water temperature was the first selected variable splitting groups of toxic MAC (14.5 °C, Fig 2). The next two selected variables were water turbidity and conductivity, with threshold values of 14.2 NTU and 0.104 mS cm^−1^, respectively. Then, intermediate conductivity values (9.16 mS cm^−1^) and high-water temperature (23.8 °C) were selected as splitting variables. Total nutrients (nitrogen and phosphorus) and wind intensity were not selected by the analysis and therefore were not relevant determining toxic ecotypes. Three ecotypes were identified in water with temperature between 14.0 °C and 22.4 °C.

As shown in Table 1, the identified ecotypes differed in their environmental preferences. Ecotype A was present in brackish waters at temperatures higher than 14.5 °C and more than 14.2 turbidity units, while ecotype B was found under the same turbidity conditions than A but in freshwater and at high water temperature (> 23.8 °C, Table 1). Ecotype C was the most frequently found, it preferred warm freshwaters and had a high amount of toxic potential (number of *mcyJ* copies per mL). On the other hand, Ecotype D inhabits at water temperature >14.5 °C but has a wide range of conductivity preference. Finally, ecotypes E and F dwells in cold water (water temperature < 14.5 ºC) but slightly differ in their conductivity preferences (0.104 mScm^−1^ conductivity threshold) (Fig 2).

## 4. Discussion

Here, we tried to shed light to the mechanisms shaping cyanobacterial diversity by analysing communities of toxic cyanobacteria inhabiting a wide array of environmental conditions that can impose hard conditions for their survival (e.g. salinity). The study deals with complex communities of organisms genetically difficult to distinguish using common molecular marker genes (Kato, 1991; Otsuka et al., 2001). Thus, we investigated if the worldwide success of toxic MAC organisms is due to the existence of ecotypes having different environmental preferences. Our approach was a combination of a molecular methods for genotyping (HRMA of *mcyJ* gene amplicons) with machine learning techniques (*f*CART) that allowed us to detect groups of MAC toxic genotypes with distinctive ecological niches, delimiting MAC ecotypes. This study complement and expand the scope of previous works using HRMA to analyse the diversity of natural microbial communities (based on the 16S rDNA gene; (Hjelmsø et al., 2014; Kim and Lee, 2014; Zeyoudi et al., 2015) by incorporating new genetic regions as diversity markers and using machine learning tools. The *f*CART proved to be a useful technique to classify toxic MAC genotypes based on their HRMA melting profiles by selecting the environmental variables with a high discriminatory power. An advantage of using an HRMA-based method instead of amplicon sequencing and further analysis of the reads to explore sequence diversity relies in the fact that, although both are based on PCR of a target gene, the former is faster (real-time data acquisition), economic and does not involve bioinformatic analysis (Vossen et al., 2009).

Microbial species can be seen as genetic, phenotypic and ecologically similar units and the task of microbial ecologists is to understand how these units are originated and selectively optimized to coexist occupying different niches or, conversely, how much these units overlap not only genetically but also ecologically (Shapiro and Polz, 2014). In the case of MAC organisms, Komárek, (2016) proposed that the diversity of this genus has not yet been solved owing to difficulties to morphological differentiation and lack of resolution in phylogenetic markers, such as 16S rRNA gene. This prompted to explore for phylogenetically informative genes that should be conducted in order to detect ecologically coherent genotypes. In this sense, Rinta-Kanto and Wilhelm (2006) studied the genetic diversity of potentially toxic *Microcystis* based on the *McyA* amino acid sequence and reported new sequences of the *mcyA* gene (Rinta-Kanto and Wilhelm, 2006). Using *mcyA* amplification and denaturing gradient gel electrophoresis (DGGE), Hu et al. (2016) found that the microcystin variants were related to the band pattern obtained, implying that the composition of *Microcystis* community determined the kind of microcystin produced (Hu et al., 2016).

It has been shown that *mcyA* has several recombination regions (Tanabe et al., 2004) and multiple recombination events were detected within the N-methyltransferase -domain of *mcyA* and the adenylation-domain of *mcyB* and *mcyC* sequences, suggesting that recombination within and between *mcy* genes contributes to their genetic diversity. However, no recombination was detected in *mcyJ* sequences, with a single copy in toxic strains (Tanabe et al., 2009), and conserved enough to address the variability of toxic populations. Taking this into account, the approach applied in the present work is a novel way to gain insight into the MAC diversity and to elucidate the mechanisms underlying the worldwide success of this bloom-forming cyanobacterium.

Variability in physical and hydrological variables (temperature, salinity and turbidity) were relevant defining MAC community structure while nutrients concentration were not, suggesting a prominent role of local conditions in spite of trophic state, due to the high availability of nutrients at all sites and sampling times (TP ~60 μg L^−1^, TN~0.9 mg L^−1^) (Figure 2, Table 1). These variables are directly related to the physical aspects that characterize ecosystem dynamics in space and time. For example, turbidity and conductivity defines the estuarine portion of the system while temperature defines seasonal changes. The finding of ecotypes representative of reservoir, riverine or estuarine ecosystems is in agreement with the ability of MAC to proliferate in a wide range of environmental scenarios, including brackish waters, when nutrients concentration is elevated by anthropogenic eutrophication. This is consistent with the worldwide distribution of MAC and its current proliferation and has been found for other cyanobacteria, such as *Prochlorococcus* and *Cylindrospermopsis* (Moore, 1998; *Piccini et al., 2011*). In the case of *Prochlorococcus*, at least six phylogenetic clades differing in physiology and occupying distinct niches in the ocean have been found, which differentiation is proposed to be based on light intensity, temperature and nutrients (Malmstrom et al., 2013; Martiny et al., 2009; Moore, 1998). Different ecotypes of *C. raciborskii* have been described based also on phylogenetic markers, morphology, tolerance to different light intensities, affinities to low or high phosphate concentrations and toxicity (Dokulil, 2000; Piccini et al., 2011). Altogether, the findings suggest that the generation of ecotypes might be a common cyanobacterial strategy to proliferate and succeed.

A study in Lake Erie (USA) found that relative abundances of some genotypes changed temporally, indicating the existence of different genotypes adapted to particular environmental characteristics (Hu et al., 2016). In the same ecosystem, Berry et al. (2017) analysed *Microcystis* oligotypes using a computational method (oligotyping) based on the V4 region of the 16S rRNA gene and found changes in *Microcystis* oligotypes composition related to spatial nutrient gradients. A similar research in Lake Taihu (China) found a clear temporal but not spatial distribution of toxic genotypes composition using DGGE of the *mcyJ* gene (Wang et al., 2012), being nitrate the variable explaining the variation in toxic genotype composition. Moreover, Tromas et al. (2018) reported that niche separation within *Microcystis* genus occurred mainly related to total dissolved nitrogen preferences and temperature. These results suggest that the presence of toxic ecotypes of MAC in those ecosystems would be associated to nutrients regime. However, in eutrophic or hypertrophic ecosystems, as observed here, nutrients are no longer shaping genotype richness. In this sense, Kim et al. (2010) found that *mcyJ* gene diversity assessed by DGGE was reduced during summer compared with spring and autumn, pointing to an effect of water temperature in the selection of different toxic populations (Kim et al., 2010).

The ecotypes of toxic MAC that were associated with brackish water (A and E) showed low concentration of *mcyJ*, meaning that in the estuary toxic MAC are rare. These results support our working hypothesis and strengthen previous information obtained from the same ecosystem, which showed a general decrease of MAC biomass and abundance of *mcy*-harbouring cells from fresh-to marine water (Kruk et al., 2017; Martínez de la Escalera et al., 2017). It has been reported that salinity concentration above the organisms’ optimum conditions causes osmotic stress, decreases photosynthesis rates and might induce cell lysis (Chen et al., 2015; Orr et al., 2004; Sabart et al., 2009; Zhang et al., 2010), leading to a decrease in abundance and biomass and precluding their detection by classical microscopy counts (Segura et al., 2017). In addition, recent studies demonstrated that some *M. aeruginosa* strains acquired the ability to produce an osmoprotectant (such as sucrose) by horizontal gene transfer, generating salt-tolerant genotypes (Tanabe et al., 2018). Thus, toxic MAC ecotypes from the estuary would be composed by slow growing, salt-tolerant organisms.

Several models have been proposed to explain the origin of bacterial ecotypes (Cohan, 2011). Among them, the Stable Ecotype model define stable and long-standing ecotypes where periodic selection limits the diversity, while the Species-Less model assumes rapid invention and extinction of ecotypes and little periodic selection, especially under environmental conditions that are rapidly changing. In Species-Less model, ecotypes evolve to invade a new ecological niche or where an environment undergoes a succession process and organisms at a site must adapt to rapidly changing conditions (Kopac and Cohan, 2011), such as those found at the assessed environmental gradient. Nonetheless, further studies should be performed in order to determine the actual speciation model taking place for MAC toxic ecotypes. Based on the high variability environmental dynamics of the addressed gradient system (Uruguay river - Río de la Plata estuary), the more suitable model to explain the origin of MAC ecotypes adapted to estuarine conditions would be Species-Less, although further work explicitly addressing this hypothesis must be carried out.

Sabart et al. (2009) studied spatial-temporal changes of *Microcystis* diversity in inter-connected freshwater ecosystems (reservoirs, ponds and river) based on the internal transcribed spacer (ribosomal ITS). They found that *Microcystis* populations were genetically different over short distances (~ 20 km) and that populations observed in the main reservoir were different from those found downstream, suggesting that they had different niches. However, at larger, global geographical scale the connections between phylogenetic relationships and genetic structure of *Microcystis* communities and environmental conditions was not detected (van Gremberghe et al., 2011). A possible explanation is that fast-evolving molecules, such as ribosomal ITS, can exhibit high levels of homoplasy, which increases the noise in the phylogenetic signal. Here, some ecotypes were detected in sites as far as ~500 km away, revealing that local environmental conditions are more relevant than distance.

In sum, we found ecotypes that are composed by a group of closely-related *Microcystis* organisms (perhaps related morpho-species), sharing ecological and toxicity characteristics. The existence of six ecotypes of toxic MAC associated to different environmental settings through the wide environmental gradient assessed lead us to hypothesize that the reservoir, which displays high MAC biomasses and diversity through the whole year, might act as a source or seed bank of toxic ecotypes. As MAC toxic organisms are transported downstream through the river and into Río de la Plata estuary, populations are selected according to their fitness in the encountered conditions, giving birth to different ecotypes. As far as we know, this is the first study combining a highly sensitive molecular technique (HRMA) with functional data analysis (*f*CART) to detect ecotypes and hypothesize the main mechanisms driving their selection. Temperature, conductivity and turbidity were the main environmental variables driving toxic genotype diversity and modulating ecotypes occurrence and distribution in Uruguay river and Río de la Plata.

## Acknowledgements

We thank Asociación Honoraria de Salvamentos Marítimos y Fluviales (ADES) and Comisión Técnico-Mixta de Salto Grande (CTM-Salto Grande) for their valuable help to perform samplings. This study was supported by grant ANII-ALGAS (Laboratorio Tecnológico del Uruguay), by PEDECIBA-Biología and by the Agencia Nacional de Investigación e Innovación of Uruguay (ANII) and Proyecto ECOS (Aprendizaje Estadístico para la Modelización y Análisis de Recursos Naturales). We thank Dr. Frederick Cohan for her critical and helpful review of the manuscript. This work was carried out in partial fulfilment of the requirements of G. Martínez de la Escalera for the doctoral degree from PEDECIBA (University of Uruguay).

